# The sulfated peptide CLEL6 is a negative regulator of anthocyanin biosynthesis in *Arabidopsis thaliana*

**DOI:** 10.1101/2022.11.23.517704

**Authors:** Eric Bühler, Elisa Fahrbach, Andreas Schaller, Nils Stührwohldt

**Affiliations:** Department of Plant Physiology and Biochemistry, Institute of Biology, University of Hohenheim, 70593 Stuttgart, Germany

**Keywords:** TPST, peptide signaling, tyrosine sulfation, anthocyanins, CLEL6, light stress

## Abstract

Post-translationally modified peptides are now recognized as important regulators of plant stress responses. Here we identified the small sulfated CLE-LIKE6 (CLEL6) peptide as a negative regulator of stress-induced anthocyanin biosynthesis. The expression of *CLEL6* and its negative effect on anthocyanin biosynthesis were strongly down-regulated by light. The function of CLEL6 depends on proteolytic processing of the CLEL6 precursor by the subtilisin-like serine proteinase 6.1 (SBT6.1), and on tyrosine sulfation by tyrosylprotein sulfotransferase (TPST). Loss of function mutants of either *sbt6.1* or *tpst* showed significantly higher anthocyanin accumulation upon light stress. The overaccumulation phenotype of *sbt6.1* and *tpst* was suppressed by application of mature CLEL6. Further confirming the role of CLEL6 as an inhibitor of anthocyanin biosynthesis, overexpression and external application of CLEL6 inhibited the expression of genes involved in anthocyanin biosynthesis in etiolated and light-stressed seedlings. Small post-translationally modified peptides are known to be perceived by leucine-rich-repeat receptor like kinases. Through a genetic approach, using a ROOT MERISTEM GROWTH FACTOR 1 INSENSITIVE (RGI) receptor quintuple mutant, we could show the essential function of the RGI receptor family in CLEL6 signaling. Our data indicate that CLEL6 inhibits anthocyanin biosynthesis through RGI receptors in dark-grown seedlings, and that this inhibition is released when CLEL6 expression is down-regulated upon transition to light.

**One sentence summary:** The formation of CLEL6 as a negative regulator of anthocyanin biosynthesis depends on proteolytic processing by SBT6.1, post-translational modification by TPST, and perception by RGI receptors.

## Introduction

The apical hook, hypocotyl elongation, and closed cotyledons are hallmarks of skotomorphogenic development of dark-grown seedlings struggling to reach the light. Skotomorphogenesis relies on a group of basic helix-loop-helix transcriptional repressors named PHYTOCHROME-INTERACTING FACTORs (PIFs) that inhibit the expression photomorphogenic genes like *HEMA1, GUN5, LHCA1* and *PS-AE1* (Duek and Fankhauser, 2005; Monte et al., 2007; Shin et al., 2009). In addition, positive regulators of photomorphogenesis, the bZIP transcription factors ELONGATED HYPOCOTYL 5 (HY5) and LONG HYPOCOTYL IN FAR-RED (HFR1), are degraded by the ubiquitin ligase complex CONSTITUTIVE PHOTOMORPHOGENESIS 1-SUPPRESSOR OF PHYA-105 (COP1-SPA) in the dark (Osterlund et al., 2000; Jang et al., 2005). Upon perception of light, phytochrome photoreceptors are activated leading to the degradation of PIFs and re-organization of the COP1-SPA complex. Thereby, photomorphogenic genes are induced, stem elongation comes to a halt, the apical hook opens, cotyledons expand, functional chloroplasts are formed, and photosynthesis ensues (Leivar et al., 2008; Jiao et al., 2007; Sheerin et al., 2015).

When chloroplast biogenesis is disturbed, or when chloroplasts are dysfunctional as a result of environmental stress, GENOMES UNCOUPLED (GUN1) mediates retrograde plastid-to-nucleus signaling to inhibit the expression of photomorphogenic genes and to prevent photo-oxidative damage (Koussevitzky et al., 2007). GUN1-mediated retrograde signaling also controls the accumulation of anti-oxidative anthocyanins that protect plants from stress-induced reactive oxygen species (ROS) (Richter et al., 2020; Nakabayashi et al., 2014; Shao et al., 2007). The genes coding for the enzymes in the anthocyanin biosynthetic pathway are grouped into early biosynthesis genes including *CHALCONE SYNTHASE* (CHS), *CHALCONE ISOMERASE* (CHI), and *FLAVANONE 3-HYDROXYLASE* (F3H), and late biosynthesis genes including *DIHYDROFLAVONOL 4-REDUCTASE* (DFR), *LEUCOANTHOCYANIDIN OXYGENASE* (LDOX) and *UDP-GLUCOSE:FLAVONOID 3-O-GLUCOSYLTRANSFERASE* (UF3GT). Expression of these genes is controlled by numerous R2R3 MYB transcription factors. This includes PRODUCTION OF ANTHOCYANIN PIGMENT1 (PAP1) for the regulation of late biosynthesis genes (Xie et al., 2006). Early biosynthesis genes are regulated by another group of R2R3-MYB transcription factors including MYB11, MYB12, MYB111 (Stracke et al., 2007). The GOLDEN-2-LIKE1 (GLK1) transcription factor mediates anthocyanin accumulation in response to sucrose (Zhao et al., 2021). In contrast to these positive regulators, and consistent with their relevance for light stress protection, anthocyanin biosynthetic genes are suppressed in dark-grown seedlings by PIF3 and HY5 that bind directly to the promoters of *CHS, CHI, F3H* and *LDOX*(Shin et al., 2007).

The regulation of plant stress responses also involves post-translationally modified peptides, particularly those containing a sulfated tyrosine residue. For example, PHYTOSULFOKINE (PSK) contributes to drought stress tolerance in Arabidopsis (Stührwohldt et al., 2021). In tomato, PSK triggers drought-induced abscission of flowers and fruits (Reichardt et al., 2020). Immune responses in Arabidopsis and tomato are also regulated by PSK signaling ((Mosher et al., 2013; Zhang et al., 2018), Igarashi et al., 2012). Likewise, CLE-LIKE9 (CLEL9) signaling enhances resistance against bacterial pathogens, and CLEL4 triggers innate immune responses in Arabidopsis (Stegmann et al., 2022)(Wang et al., 2021). PLANT PEPTIDE CONTAINING SULFATED TYROSINE (PSY) peptides have been implicated in abscisic acid-mediated stress responses and immune signaling (Shen et al 2013)(Tost et al., 2021), and unidentified sulfated peptides are involved in the response to phosphate deficiency (Kang et al., 2014).

The formation of these peptides relies on subtilisin-like serine proteinases (subtilases, SBTs) for processing (Schaller et al., 2018), and on tyrosylprotein sulfotransferase (TPST) for post-translational sulfation (Komori et al., 2009). In case of CLEL peptides, processing by SBT6.1 is required for maturation and secretion, and sulfation by TPST for bioactivity (Stührwohldt et al., 2020; Ghorbani et al., 2016). Bioactivity of PSK depends on processing by SBT3.8 and Phyt2 in tomato and Arabidopsis, respectively (Stührwohldt et al., 2021; Reichardt et al., 2020), and on TPST for receptor recognition (Wang et al., 2015). In case of TWISTED SEED1 (TWS1), tyrosine sulfation is required for both, receptor binding and processing by SBT1.8 (Royek et al., 2022; Doll et al., 2020; Stintzi and Schaller, 2022)

In this work we show that anthocyanin accumulation in Arabidopsis is regulated by CLEL6. We further show that this regulatory function of CLEL6 depends on proteolytic processing by SBT6.1, tyrosine sulfation by TPST, and perception by RGF1 INSENSITIVE (RGI) receptors.

## Results

### Photomorphogenesis and anthocyanin biosynthesis are mis-regulated in *tpst-1*

Since PIFs are negative regulators of photomorphogenesis, the *pifq* quadruple mutant lacking PIF1, PIF3, PIF4 and PIF5 has a constitutive photomorphogenic (*cop*)-like phenotype in the dark (Leivar et al., 2008). We observed that etiolated *tpst-1* seedlings display short hypocotyls and open cotyledons thus phenocopying the *pifq* mutant (Figure 1, A-C). The *cop*-like phenotype of *pifq* is suppressed when retrograde signaling is activated by lincomycin (linc), a drug that inhibits protein biosynthesis in plastids and thus interferes with chloroplast development (Martín et al., 2016). Therefore, to confirm the phenotype of *tpst-1* as *cop*-like, we treated etiolated seedlings with linc and observed that early opening of cotyledons was indeed suppressed (Figure 1D). In light-grown seedlings, however, the activation of retrograde signaling by linc led to accumulation of photo-protective anthocyanins, and anthocyanin levels were significantly higher in the *tpst-1* mutant compared to the wild type (Figure 1E). Surprisingly, *tpst-1* seedlings accumulated more anthocyanins than the wild type even in the absence of linc (Figure 1E) suggesting that this aspect of the *tpst-1* phenotype is independent of retrograde signaling. The data indicate that TPST activity is required to suppress photomorphogenesis in the dark and to control the accumulation of anthocyanins. In this study, we focused on the TPST-dependent regulation of anthocyanin biosynthesis. Since TPST is required for the post-translational modification and bioactivity of signaling peptides, we hypothesize that a sulfated peptide is involved in this process.

**Figure 1:**
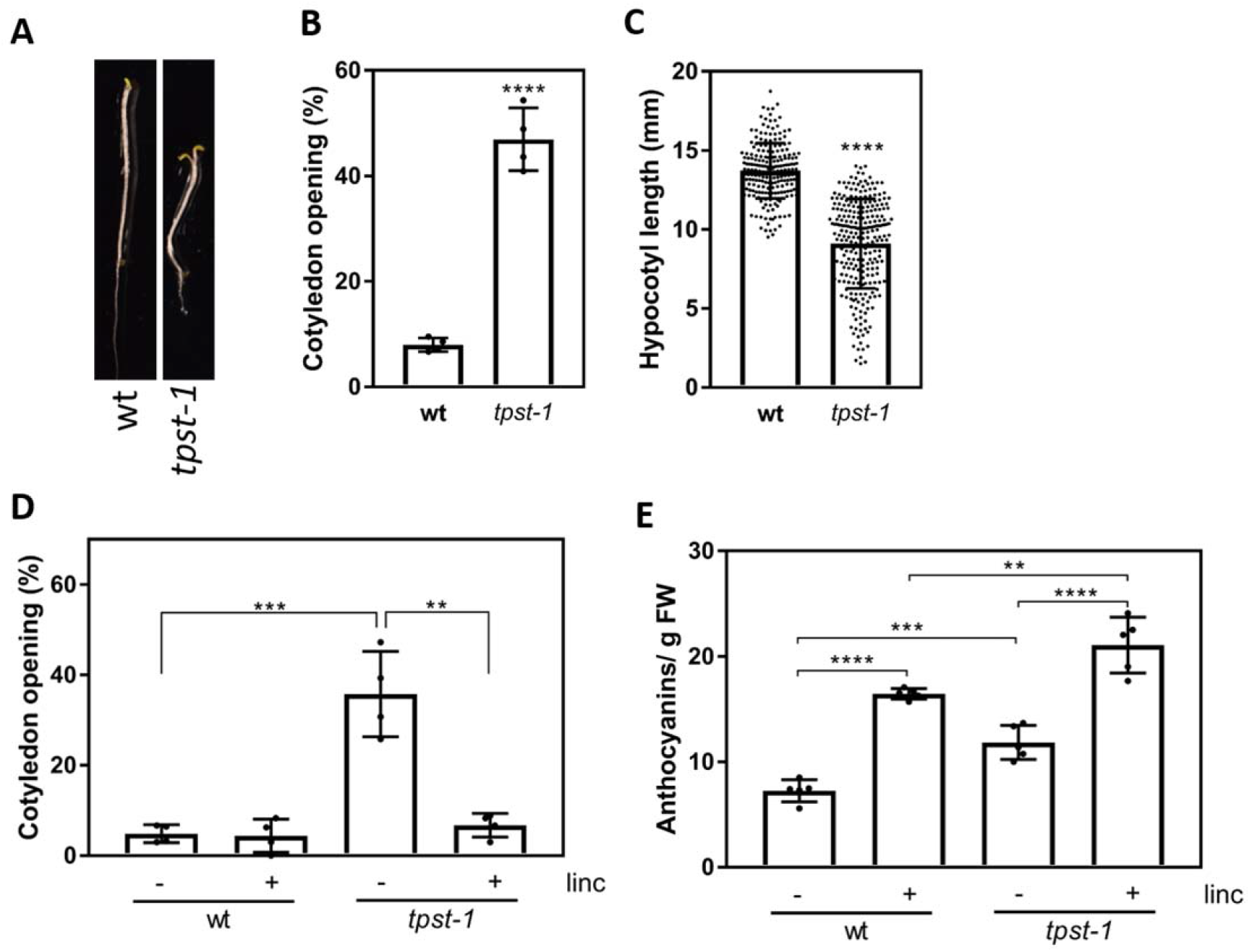
*tpst-1* displays an early photomorphogenesis phenotype and accumulates high amounts of anthocyanins. **A** Representative five-day-old etiolated wild-type (wt) and *tpst-1* seedlings. **B, C** Quantification of **(B)** open cotyledons (in %) and **(C)** hypocotyl length (mm) of five-day-old etiolated wt and *tpst-1* seedlings. **D** Fraction of plants with open cotyledons in five-day-old etiolated wt and *tpst-1* seedlings. Lincomycin (linc, 0.5 mM) was added to the media as indicated (-/+). **E** Quantification of anthocyanin content in wt and *tpst-1* seedlings grown for seven days under short day conditions. Linc (0.5 mM) was added to the media as indicated (-/+). Data are shown as the mean ± SD. (B) n=4 independent experiment with at least 36 seedlings each. (C) n=4 independent experiment with at least 54 seedlings each. (D) n=4 independent experiment with at least 26 seedlings each. (E) n=4. Asterisks indicate significant differences (\*\**P*<0.01, \*\*\**P*<0.001, \*\*\*\**P*<0.0001; two-tailed *t*-test).

### CLEL6 is a negative regulator of anthocyanin biosynthesis

Anthocyanin accumulation is induced by light to higher levels in *tpst-1* compared to wild-type seedlings (Figure 1E). We attribute the increase in anthocyanins in *tpst-1* to the loss of a putative signaling peptide that acts as a negative regulator to restrict anthocyanins in the wild type. Therefore, to identify this negative regulator we looked for candidate peptides that are expressed at lower levels in light-treated as compared to etiolated seedlings. The expression of all known genes coding for sulfated signaling peptides was analyzed in dark-grown wild-type seedlings two hours after light exposure and compared to the etiolated control. Of all genes tested, *CLEL6* was the only one expressed at significantly (3.5-fold) lower level in light-exposed seedlings (Supplemental Figure S1). In contrast to most other *CLEL* genes that are active predominantly in the root, *CLEL6* and *CLEL9* are shoot-specific and mainly expressed in the hypocotyl and leaves, which matches the anthocyanin accumulating tissues (Fernandez et al., 2013).

CLEL6 and 9 depend on the subtilase SBT6.1 for secretion and function (Stührwohldt et al., 2020; Ghorbani et al., 2016). Like *tpst-1, sbt6.1* can thus be considered a CLEL6 loss-of-function mutant. Therefore, to confirm CLEL6 as a negative regulator of anthocyanin biosynthesis, we compared anthocyanin levels and the expression of anthocyanin biosynthesis genes in light-stressed *sbt6.1, tpst-1,* and wild-type seedlings. When five-day-old etiolated seedlings were exposed to light for 16 hours, *tpst-1* and *sbt6.1* accumulated significantly more anthocyanins than wild type (Figure 2, A and B). The accumulation of anthocyanins correlated with the expression of early (*CHS*, *F3H*) and late biosynthesis genes (*DFR, LDOX, UF3GT, UGT78D2*) and the corresponding transcription factors (*MYB11* and *PAP1*, respectively). In the *sbt6.1* mutant all tested genes were significantly upregulated (Figure 2C). In the *tpst-1* mutant *CHS*, *F3H*, *DFR*, *LDOX* and *MYB11* were also upregulated. Only for two of the late genes (*UF3GT*, *UGT78D2*) and *PAP1*, the induction in *tpst-1* was not significantly different from the wild type (Figure 1D).

**Figure 2:**
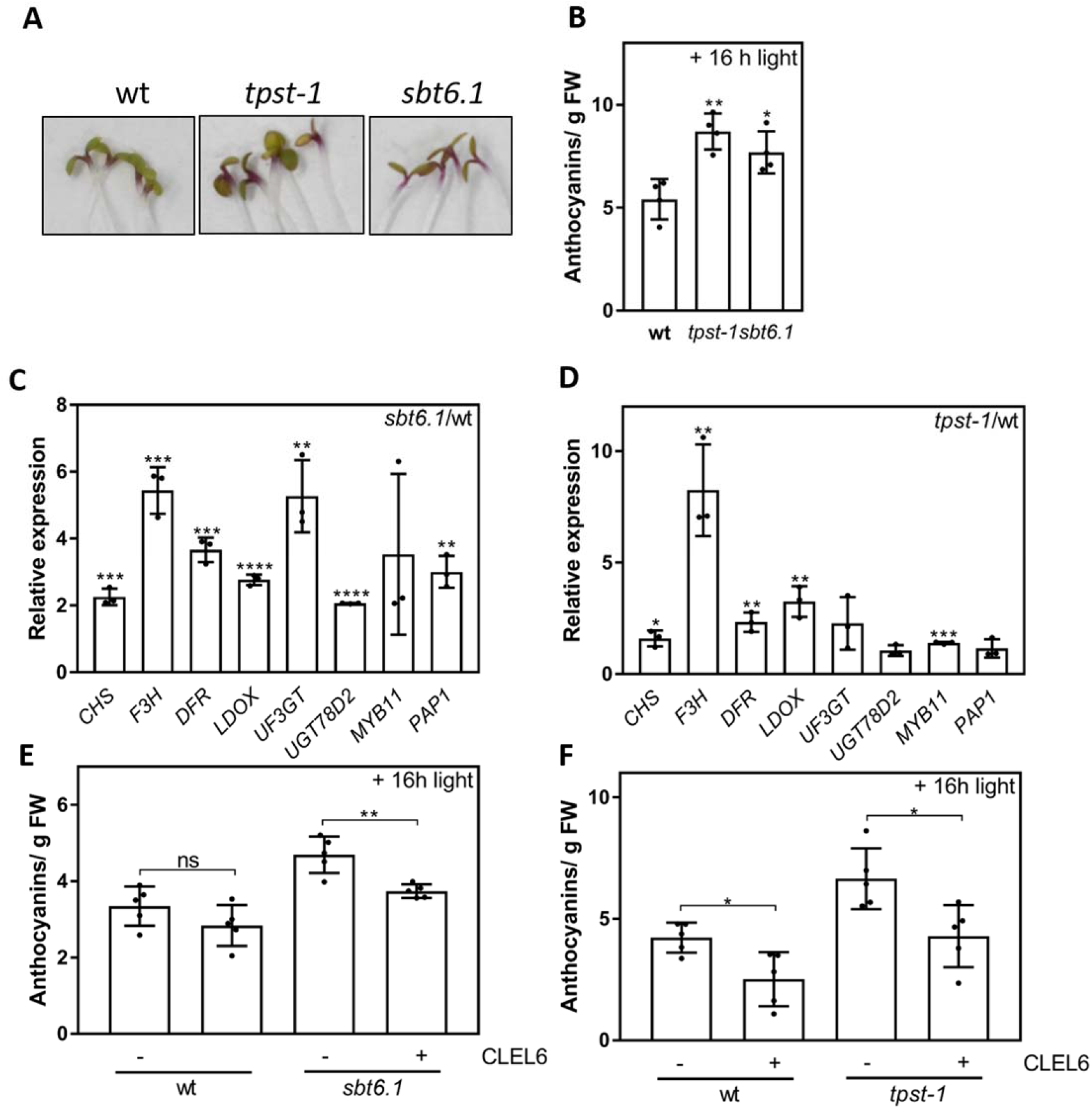
CLEL6 signaling represses anthocyanin biosynthesis. **A** Representative wild-type (wt), *tpst-1* and *sbt6.1* seedlings grown for five-days in the dark followed by 16 h in the light. **B** Quantification of anthocyanin content in five-day-old etiolated wt, *tpst-1* and *sbt6.1* seedlings after 16 h light exposure. **C, D** Relative expression of *CHS,* F3H, *DFR, LDOX, UF3GT, UGT78D2, PAP1, MYB11* in five-day-old etiolated *sbt6.1* or *tpst-1* seedlings compared to the wt control. The seedlings were exposed to light for 2 h before sampling. Gene expression was normalized to two reference genes. **E, F** Quantification of anthocyanin content of five-day-old etiolated seedlings (wt and *sbt6.1* in E; wt and *tpst-1* in F) after 16 h light exposure. 1 μM CLEL6 was added to the media as indicated (-/+) Data are shown as the mean ±SD of n=4 (B), n=3 (C, D), or n=5, (E, F) independent experiments. Asterisks indicate significant differences (\**P*<0.05, \*\**P*<0.01, \*\*\**P*<0.001; two-tailed *t*-test).

The involvement of CLEL6 in the regulation of anthocyanin biosynthesis was further substantiated by addition of the mature CLEL6 peptide (DY(SO_3_H)PQPHRKPPIHNE) to the growth medium. Addition of CLEL6 reduced anthocyanin levels in *sbt6.1* and *tpst-1* to that of the wild type (Figure 2, E and F). Confirming this result, the anthocyanin over-accumulation phenotype was also observed in a second, independent *tpst* allele (*tpst-2*, aka *hps7*; Kang et al., 2014), and complemented by treatment with CLEL6 (Supplemental Figure S2A). In contrast to CLEL6, the closely related CLEL9 peptide (DMDY(SO_3_H)NSANKKRPIHN) did not reduce anthocyanin content of the *sbt6.1* mutant (Supplemental Figure S2B). The data confirm a specific role for CLEL6 in the regulation of anthocyanin biosynthesis

### Loss of CLEL6 function results in light-independent induction of anthocyanin biosynthesis

In Arabidopsis, anthocyanins are only formed in response to light, because PAP transcription factors are destabilized by the COP1-SPA ubiquitin ligase complex in darkness (Maier et al., 2013). However, for the *tpst-1* mutant, we observed the accumulation of anthocyanins also in etiolated seedlings (Figure 3A). qPCR analysis showed that the expression of all tested anthocyanin biosynthesis genes except *MYB11* was strongly (up to 50-fold) upregulated in etiolated *tpst-1* compared to wild-type seedlings (Figure 3B). Light-independent accumulation of anthocyanins and upregulation of anthocyanin-related genes including *MYB11* was confirmed in the independent *tpst-2* allele (Supplemental Figure S3, A and B). Anthocyanin biosynthesis genes were also upregulated in the dark-grown *sbt6.1* mutant, albeit to much lower levels compared to *tpst-1*, which appeared not to be sufficient for a significant increase in anthocyanin content (Figure 3, C and D). To verify that the lack of CLEL6 is responsible for the accumulation of anthocyanins in the dark, we grew etiolated *tpst-1* seedlings on media supplemented with the mature CLEL6 or CLEL9 peptides. In contrast to CLEL9 which had no significant effect (Supplemental Figure S4A), CLEL6 treatment led to a substantial reduction in anthocyanin content of etiolated *tpst-1* seedlings (Figure 3E). Consistently, the expression level of most of the anthocyanin biosynthesis genes was much reduced in CLEL6-treated, etiolated *tpst-1* seedlings (Figure 3F). Even in the wild type, the low expression of anthocyanin-related genes in dark-grown seedlings was further reduced in presence of CLEL6 (Supplemental Figure S4B). The data indicate that CLEL6 contributes to the suppression of anthocyanin biosynthesis in the dark.

**Figure 3:**
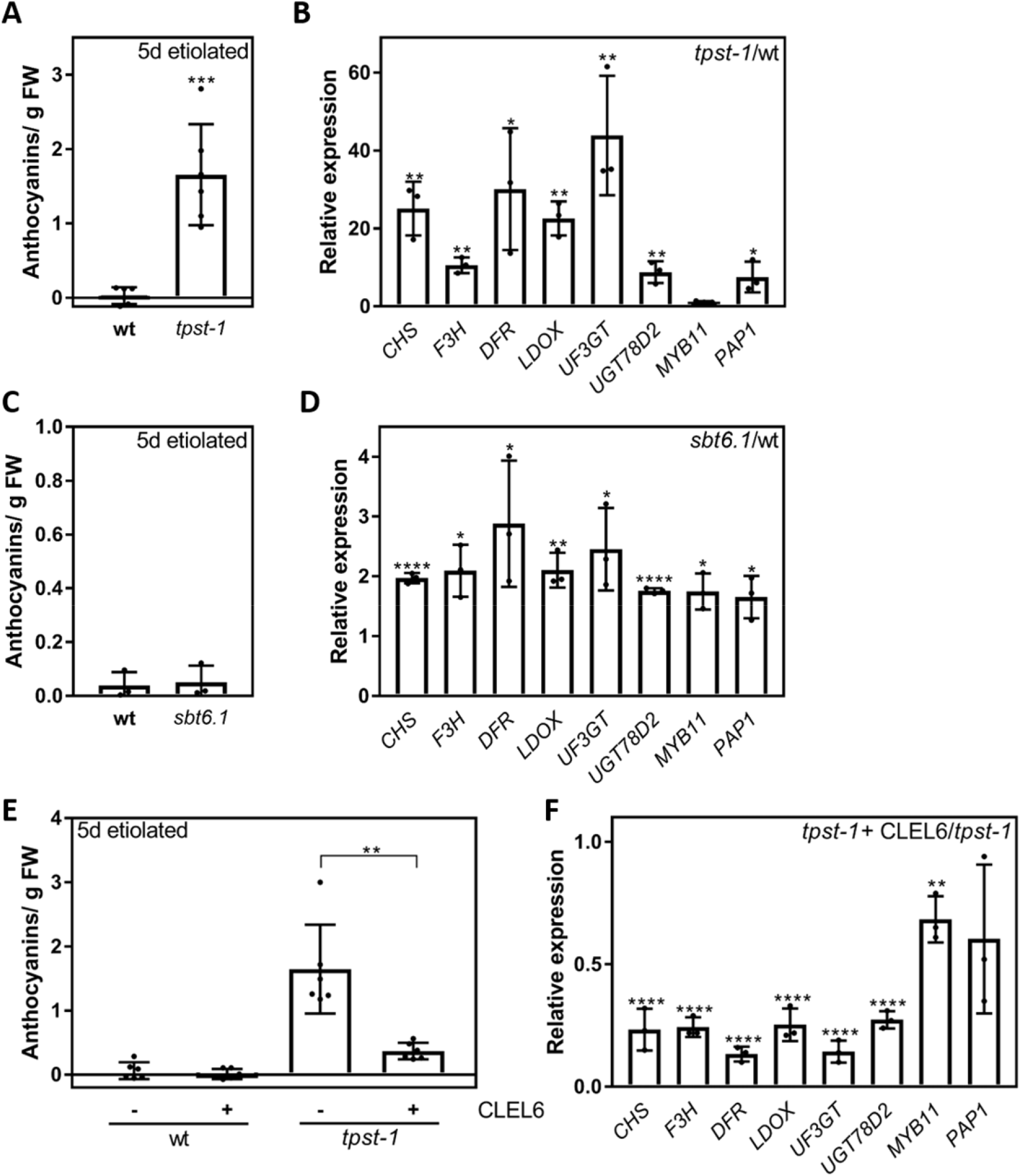
CLEL6 suppresses anthocyanin accumulation and gene expression in etiolated *tpst-1* and *sbt6.1* seedlings. **A** Quantification of the anthocyanin content of five-day-old etiolated wt and *tpst-1* seedlings. **B** Relative Expression of *CHS*, *F3H*, *DFR, LDOX, UF3GT, UGT78D2, PAP1, MYB11* in five-day-old etiolated *tpst-1* seedlings relative to the wt control. Gene expression was normalized to two reference genes. **C** Quantification of the anthocyanin content of five-day-old etiolated wt and *sbt6.1* seedlings. **D** Relative Expression of *CHS*, *F3H*, *DFR*, *LDOX, UF3GT, UGT78D2, PAP1, MYB11 of* five-day-old etiolated *sbt6.1* seedlings relative to the wt control. Gene expression was normalized to two reference genes. **E** Quantification of the anthocyanin content in five-day-old etiolated wt and *tpst-1* seedlings with 1 μM CLEL6 added in the media as indicated (-/+). **F** Relative Expression of *CHS*, *F3H*, *DFR*, *LDOX, UF3GT, UGT78D2, PAP1* and *MYB11* of five-day-old etiolated *tpst-1* seedlings grown on a media with 1 μM CLEL6 relative to the *tpst-1* control. Gene expression was normalized to two reference genes. Data are shown as the mean ±SD of x n=3 (B,C, D, F) or n=6 (A, E), biological replicates. Asterisks indicate significant differences (\**P*<0.05, \*\**P*<0.01, \*\*\**P*<0.001, \*\*\*\**P*<0.0001; two-tailed *t*-test).

### Overexpression of CLEL6 inhibits anthocyanin production

Consistent with the proposed role for CLEL6 as a negative regulator of anthocyanin biosynthesis, we observed that anthocyanin accumulation was impaired in transgenic plants constitutively overexpressing CLEL6. *35S:CLEL6* seedlings produced less anthocyanins than the wild type in response to light stress (Figure 4, A and B), and the expression of most of the anthocyanin biosynthesis genes was strongly reduced in light-stressed *35S:CLEL6* compared to wild-type seedlings (Figure 4C). In contrast, in transgenic *35S:CLEL9* seedlings, the overexpression of CLEL9 did not affect anthocyanin biosynthesis and accumulation, confirming the specific role of CLEL6 in this process (Supplemental Figure S5A). Also in dark-grown seedlings, the expression of anthocyanin biosynthesis genes was much lower in *35S:CLEL6* transgenics compared to the wild type, and formation of anthocyanins was not observed (Figure 4, D and E), while overexpression of CLEL9 (*35S:CLEL9*) had no effect on anthocyanin content (Supplemental Figure S5B). The results are consistent with those obtained in peptide complementation experiments (Figure 2, E and F, Figure 3, E and F, Supplemental Figures S2A and S4B) and confirm a specific role for CLEL6 in the downregulation of anthocyanin biosynthesis.

**Figure 4:**
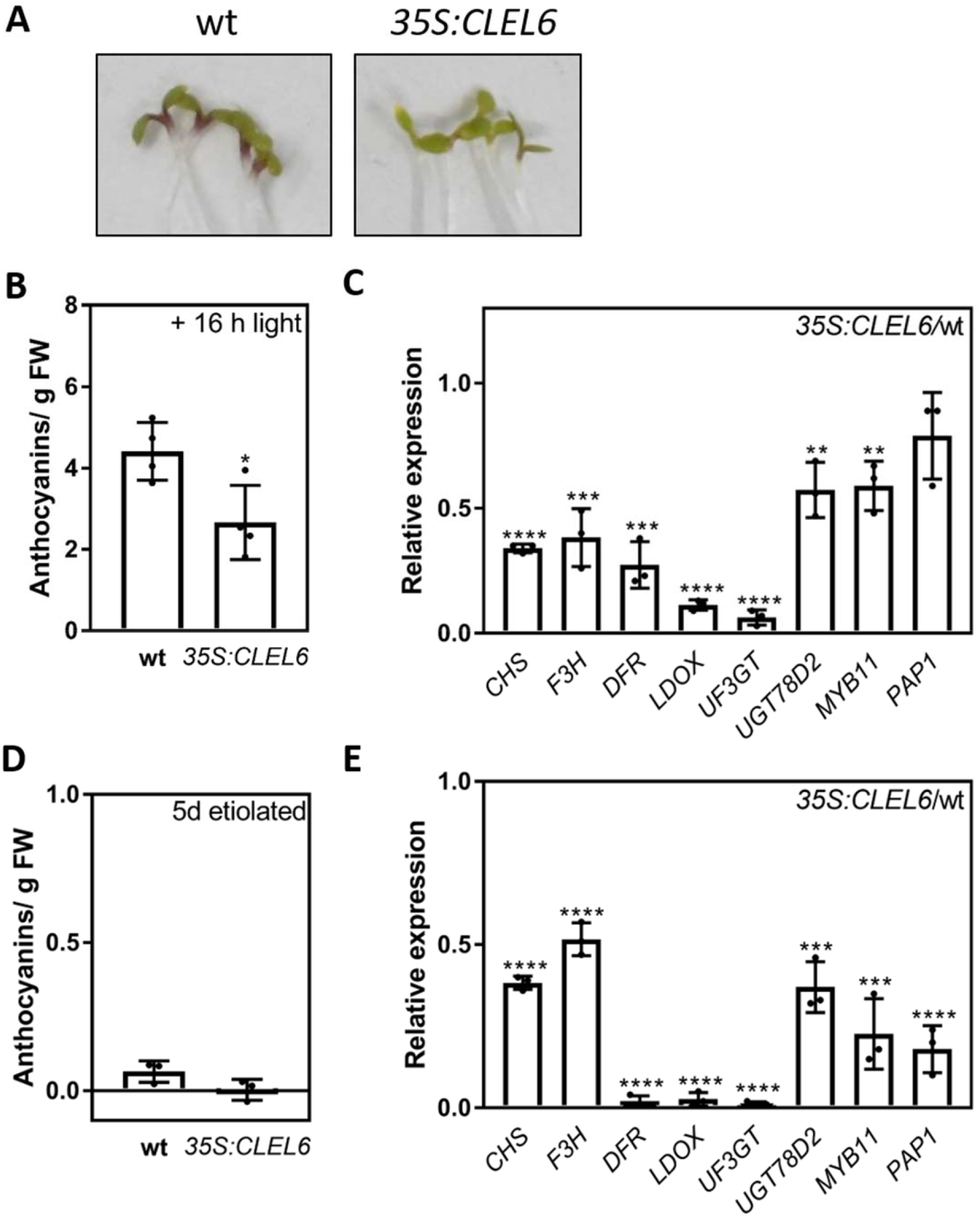
Overexpression of CLEL6 suppresses anthocyanins accumulation and biosynthesis genes. **A** Representative wt and *35S:CLEL6* seedlings grown for five-days in the dark followed by 16 h in the light. **B** Quantification of anthocyanin content of five-day-old etiolated wt and *35S:CLEL6* seedlings after 16 h exposure to light. **C** Relative Expression of *CHS*, *F3H*, *DFR*, *LDOX, UF3GT, UGT78D2, PAP1, MYB11* in five-day-old etiolated *35S:CLEL6* seedlings compared to the wt control. Seedlings were exposed to light for 2 h before sampling. Gene expression was normalized to two reference genes. **D** Quantification of anthocyanin content of five-day-old etiolated wt and *35S:CLEL6* seedlings. **E** Relative Expression of *CHS*, *F3H*, *DFR*, *LDOX, UF3GT, UGT78D2*, *PAP1*, *MYB11* of five-day-old etiolated *35S:CLEL6* seedlings relative to the wt control. Gene expression was normalized to two reference genes. Data are shown as the mean ±SE of B n=4 (B) or n=3 (C-E) independent experiments. Asterisks indicate significant differences (\**P*<0.05, \*\**P*<0.01, \*\*\**P*<0.001; \*\*\*\**P*<0.0001; two-tailed *t*-test).

### CLEL6 signaling depends on RGIs

Within the large family of leucine-rich repeat receptor kinases (LRR-RKs), RGIs (ROOT MERISTEM GROWTH FACTOR (RGF)1 INSENSITIVEs) have been identified as receptors of CLEL peptides. RGI receptors act redundantly in the perception of CLEL8 (aka RGF1) for the regulation of root meristem development (Ou et al., 2016). RGI3 perceives CLEL9 (RGF9) and is required for its immune-modulatory activity (Stegmann et al., 2022). Likewise, the activity of CLEL4 (RGF7) as a damage-associated molecular pattern relies on the RGI4 and RGI5 receptors (Wang et al., 2021). The RGI family comprises five members. While RGI1 and RGI2 are mostly expressed in the root tip (hence their name), RGI3-5 are also expressed in leaves (Song et al., 2016; Stegmann et al., 2022). To address the question whether the RGIs are important for CLEL6 signaling, we used a *rgi* quintuple mutant, *rgi5x* (Ou et al., 2016). Upon light exposure, *rgi5x* accumulated significantly more anthocyanin than the wild type, thus resembling the CLEL6 loss-of-function phenotype of the *tpst-1* mutant (Figure 5A). While anthocyanin concentration in *tpst-1* was reduced to wild-type levels when grown in presence of CLEL6, anthocyanin levels in *rgi5x* were insensitive to CLEL6 treatment (Figure 5A). Similar to the *sbt6.1* and *tpst-1* mutants that fail to produce functional CLEL6 (Figure 3, B and D), the expression of anthocyanin biosynthesis genes was elevated in *rgi5x* seedlings, in darkness as well as after light exposure (Figure 5, B and C). However, unlike *tpst-1* where we observed strong suppression of anthocyanin biosynthesis genes in response to CLEL6 (Figure 3F), CLEL6 treatment could not reduce the expression of these genes in the *rgi5x* receptor mutant (Figure 5D). CLEL6 insensitivity of *rgi5x* was also observed with respect to hypocotyl elongation. The overexpression of *CLEL6* was previously shown to cause an increase in hypocotyl length of etiolated seedlings (Ghorbani et al 2016), which we confirmed in the *35S:CLEL6* transgenic line (Supplemental Figure S6A). Consistent with this finding, hypocotyl growth was also stimulated by CLEL6 treatment in etiolated wild-type and *tpst-1* seedlings, and the effect was much reduced in the *rgi5x* mutant (Supplemental Figure S6B). The data indicate that CLEL6 activity depends on RGIs, and they confirm that CLEL6-signaling through RGI receptor(s) controls anthocyanin formation by suppression of anthocyanin biosynthesis genes (Figure 6).

**Figure 5:**
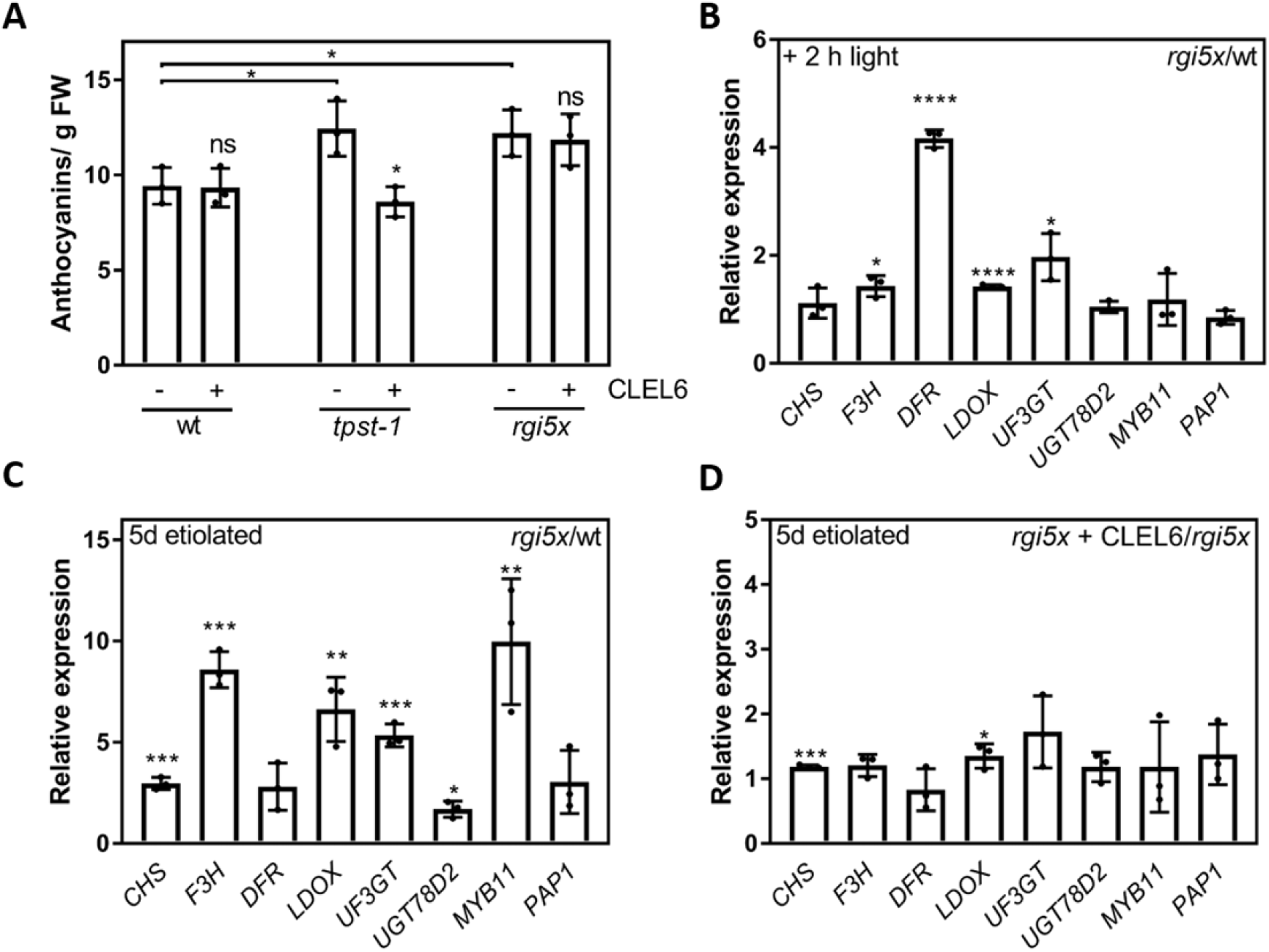
The *rgi5x* receptor mutant is insensitive to CLEL6 treatment. **A** Quantification of anthocyanin content of five-day-old etiolated wt, *tpst-1* and *rgi5x* seedlings after 16h light exposure. 1 μM CLEL6 was added to the media as indicated (−/+). **B** Relative expression of *CHS*, *F3H*, *DFR*, *LDOX, UF3GT, UGT78D2, PAP1, MYB11* in five-day-old etiolated *rgi5x* seedlings compared to the wt control. Seedlings were exposed to light for 2 h before the sampling. Gene expression was normalized to two reference genes. **C** Expression of *CHS*, *F3H*, *DFR*, *LDOX, UF3GT, UGT78D2, PAP1, MYB11* in 5-day-old etiolated *rgi5x* seedlings relative to the wt control. Gene expression was normalized to two reference genes. **D** Expression of *CHS*, *F3H*, *DFR*, *LDOX, UF3GT*, *UGT78D2*, *PAP1*, *MYB11* in 5 day-old etiolated *rgi5x* seedlings grown on a media with 1 μM CLEL6 relative to the *rgi5x* control. Gene expression was normalized to two reference genes. Data are shown as the mean ±SD for n=3 (A-D) independent experiments. Asterisks indicate significant differences (\**P*<0.05, \*\**P*<0.01, \*\*\**P*<0.001, \*\*\*\**P*<0.0001; two-tailed *t*-test).

**Figure 6:**
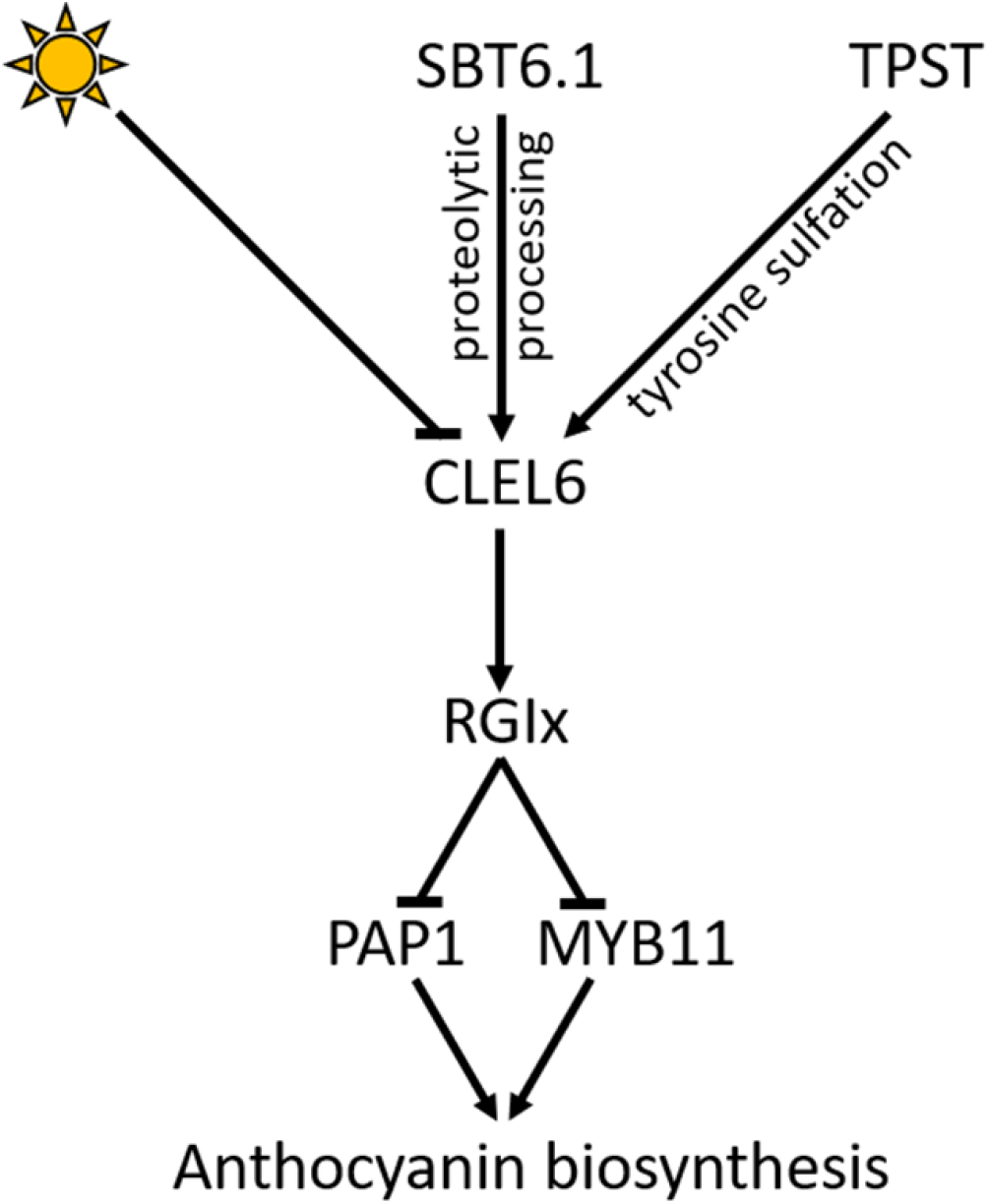
Model of CLEL6-regulated anthocyanin biosynthesis. Light downregulates the expression of *CLEL6*. CLEL6 peptide biogenesis depends on SBT6.1 for processing and on TPST for sulfation. The mature CLEL6 peptide activates RGI receptor(s). RGI-signaling downregulates the expression of the PAP1 and MYB11 transcription factors. MYB11 induces the early biosynthesis genes while PAP1 induces the late biosynthesis genes for anthocyanin formation.

## Discussion

CLEL6 is a small post-translationally sulfated peptide in the family of CLEL peptides, that are also known as ROOT GROWTH FACTORs (RGFs) or GOLVEN peptides (GLVs) (Matsuzaki et al., 2010; Meng et al., 2012; Whitford et al., 2012). These peptides are well-known for their function in root growth and development. CLELs control the size of the root meristematic zone and the stability of PLETHORA (PLT) transcription factors that are required for root stem cell maintenance (Yamada et al., 2020; Matsuzaki et al., 2010; Meng et al., 2012). CLELs are also involved in root gravitropism and have been implicated in lateral root development (Meng et al., 2012; Whitford et al., 2012). However, unlike most other CLELs that are predominantly expressed in roots, CLEL4 is induced in Arabidopsis shoots after pathogen infection (Wang et al., 2021), and CLEL6 and CLEL9 show constitutively high expression in hypocotyls and leaves (Whitford et al., 2012; Fernandez et al., 2013; Stegmann et al., 2022). Consistent with this expression pattern, CLEL6 and 9 are involved in growth and gravitropic responses of the hypocotyl (Whitford et al., 2012; Stührwohldt et al., 2020; Ghorbani et al., 2016) . The two peptides appear to act redundantly in these processes, and their formation and bioactivity depend on SBT6.1 for processing (Ghorbani et al., 2016; Stührwohldt et al., 2020) and on TPST for post-translational sulfation (Stührwohldt et al., 2020).

We report here an early photomorphogenic phenotype and mis-regulated anthocyanin biosynthesis for the *tpst* mutant, observed both in etiolated seedlings and in response to light, suggesting sulfated signaling peptides to be involved in the regulation of these processes. Mis-regulation of anthocyanin biosynthesis was also observed for the *sbt6.1* mutant that is impaired specifically in the biogenesis of the CLEL peptides. Confirming a role for CLEL peptides in the regulation of anthocyanin biosynthesis, the overaccumulation phenotype of *tpst* and *sbt6.1* mutants was suppressed by treatment with CLEL6. Furthermore, CLEL6 inhibited the expression of anthocyanin biosynthesis genes in etiolated as well as light-exposed seedlings, and its activity was lost in the *rgi5x* receptor mutant. However, unlike hypocotyl growth and gravitropic response which are redundantly controlled by CLEL6 and 9, CLEL9 treatment, or overexpression of the CLEL9 peptide precursor, did not affect anthocyanin biosynthesis and accumulation. The data indicate that the inhibition of anthocyanin biosynthesis is mediated specifically by CLEL6. Consistent with the proposed role for CLEL6 as a negative regulator of anthocyanin biosynthesis during skotomorphogenesis, *CLEL6* was the only one of all sulfated peptide precursor genes that is expressed in dark-grown seedlings and downregulated by light.

We propose that anthocyanin biosynthesis is inhibited by CLEL6 during skotomorphogenesis, and this inhibition is released when the expression of *CLEL6* is downregulated after light exposure (Figure 6, Supplemental Figure S1). Since light-dependent formation of anthocyanins is also controlled by transcription factors that act as negative (PIFs) and positive (HY5) regulators of photomorphogenesis (Shin et al., 2007; Lee et al., 2007), we hypothesize that they may do so by regulating *CLEL6* expression. HY5 activates photomorphogenic genes including those for anthocyanin biosynthesis (Lee et al., 2007). HY5 also induces the expression of *PAP1* which in turn stimulates late anthocyanin biosynthesis genes (Shin et al., 2013). Repression of *CLEL6* expression in the light may constitute an additional mechanism by which positive regulators like HY5 could contribute to light-dependent anthocyanin formation.

Alternatively, expression of *CLEL6* in etiolated seedlings and its downregulation after light exposure may be mediated by transcription factors like PIFs, that are inactivated by light. The PIF family comprises eight members and they are key regulators in the transition from skotomorphogenic development to photomorphogenesis. PIFs accumulate in darkness and are rapidly degraded in response to light (Duek and Fankhauser, 2005; Leivar et al., 2008). Most important for the shift from skoto- to photomorphogenesis are PIF1 and PIF3 (Shin et al., 2009; Shen et al., 2005). CLEL6 activity in etiolated seedlings matches that of PIFs. Like PIFs, CLEL6 promotes hypocotyl growth (Ghorbani et al., 2016)Supplemental Figure S6) and inhibits anthocyanin biosynthesis (Figures 3 and 4), and these activities are reduced when seedlings are exposed to light. CLEL6 may thus act downstream of PIFs, thereby contributing to the transition from skoto- to photomorphogenesis.

Evidence supporting the hypothesis that *CLEL6* expression is controlled by PIFs was found in published sequence data. Reduced expression of *CLEL6* in the *pifq* mutant compared to the wild type was reported in three independent RNA-seq studies (Martín et al., 2016; Oh et al., 2012; Richter et al., 2010). Two further studies reported lower expression of *CLEL6* in a *pif4pif5* double mutant compared to wild type (Lorrain et al., 2008; Nozue et al., 2011). In a promoter analysis by Martín et al. (2016), *CLEL6* was predicted as a direct target of PIF1, and Ou et al. (2016) reported that PIF4 binds to the promoters of *CLEL6* and *TPST*. Taken together, these data provide strong evidence for the hypothesis that the expression of *CLEL6* in etiolated seedlings and its downregulation upon light exposure are controlled by the PIF transcription factors. The negative effect of CLEL6 on the expression of the *MYB11* and *PAP1* transcription factors and of the anthocyanin biosynthesis genes is thus released when PIFs are degraded, allowing for the synthesis and accumulation of photoprotective pigments in response to light.

## METHODS

### Plant materials and growth conditions

Plants were grown on 0.5 × Murashige and Skoog medium (Murashige and Skoog, 1962) supplemented with 1% sucrose and 0.38% gelrite. One μM CLEL6 (DY(SO3H)PQPHRKPPIHNE; Pepscan, Lelystad, The Netherlands), 1 μM CLEL9 (DMDY(SO_3_H)NSANKKRPIHN; SciLight Peptide, Beijing, China) or 0.5 mM lincomycin (Sigma, St. Louis, Missouri, USA) were added as indicated. Arabidopsis seeds were sterilized for 15 minutes in 70% ethanol. Seeds were stratified for 2 days at 4°C. Germination was induced by a 4h-6h light treatment (120 μmol m^−2^ s^−1^ white light). Afterwards, plants were grown at 22 °C in darkness or under short-day (12h/12h) conditions. For light stress experiments, seedlings were grown for five days in the dark and transferred into constant light (120 μmol m^−2^ s^−1^ white light) for 2 or 16 hours as indicated.

### Arabidopsis lines

All transgenic lines and mutants used for this study were described previously; *tpst-1 (Komori et al., 2009), tpst-2 (Kang et al., 2014), sbt6.1 (Rautengarten et al., 2005), 35S:CLEL6* and *35S:CLEL9 (Whitford et al., 2012), and rgi5x* (Ou et al., 2016).

### Quantification of hypocotyl length and cotyledon opening

Seedlings were grown for five-day in darkness. The cotyledons were scored as open when they were separated from each other. Hypocotyl length was determined using Image J.

### Anthocyanin extraction and measurement

The protocol was adapted from (Nakata et al., 2013). Approximately 30 seedlings were harvested for each sample and fresh weight was determined. Samples were frozen in liquid nitrogen. Frozen samples were homogenized in a TissueLyser LT (Qiagen, Venlo, Netherlands). 200 μl of extraction buffer (45% methanol, 5% acetic acid) were added and samples were vortexed vigorously. The cell debris was removed by centrifugation at 13000 rpm in a microfuge at 4°C for 10 minutes. The supernatant was transferred into a new microfuge tube. Centrifugation and transfer into a new tube were repeated. The Absorption was measured at 530 nm and 657 nm in a TECAN Spark plate reader (Tecan Deutschland GmbH, Crailsheim, Germany). Anthocyanin content was calculated as ([Abs530 - (0.25 × Abs657)] × 2) / g FW.

### qPCR

Total RNA was extracted using TRIzol (ABP Biosciences, Beltsville, Maryland, USA) following the manufacturers protocol. Up to 5 μg RNA was used for cDNA synthesis by RevertAid reverse transcriptase (Thermo Fisher Scientific, Waltham, Massachusetts, USA) using oligo dT Primers. qPCR Primers are listed in Supplemental Table S1. qPCR was performed in a CFX96 Real-Time PCR Detection system (Bio-Rad, Hercules, California, USA). Experiments were performed in three biological and two technical replicates. Expression data were normalized against the two housekeeping genes *ACTIN2* (*At3g18780*) and *TUB4* (*At5g44340*).

## Acknowledgements

Authors thank Lhana Stein for excellent technical support. We also thank Prof. Liu for the *tpst-2* line, Ana Fernandez for the *35S:CLEL6* and *35S:CLEL9* line *and* Prof. Li for the *rgi5x* line.

## Supplemental Figures

**Supplemental Figure S1: *CLEL6* expression is downregulated by light.** Expression of the genes coding for sulfated peptides in five-day-old etiolated wild-type wt seedlings after 2h of light exposure relative to the dark control. Gene expression was normalized to two reference genes. Data are shown as the mean ±SD for n=3 independent experiments. Asterisks indicate significant differences (\**P*<0.05, \*\**P*<0.01, \*\*\*\**P*<0.0001; two-tailed *t*-test).

**Supplemental Figure S2. CLEL6 complements the anthocyanin accumulation phenotype, CLEL9 does not. A** Quantification of anthocyanin content in five-day-old etiolated wild-type (wt) and *tpst-2* seedlings after 16 h of light exposure. 1 μM CLEL6 was added as indicated (−/+). **B** Quantification of anthocyanin content in five-day-old wt and *sbt6.1* seedlings 16 h after exposure to light 1 μM CLEL9 was added as indicated (−/+). (A,B) Data are shown as the mean ±SD for n=3 independent experiments. Asterisks indicate significant differences (\**P*<0.05; two-tailed *t*-test).

**Supplemental Figure S3. Anthocyanin accumulation and biosynthesis genes are upregulated in the independent *tpst-2* mutant. A** Quantification of anthocyanin content in five-day-old etiolated wild-type (wt) and *tpst-2* seedlings. **B** Expression of *CHS*, *F3H*, *DFR*, *LDOX, UF3GT, UGT78D2, PAP1, MYB11* in five-day-old etiolated *tpst-2* seedlings relative to the wt control. Gene expression was normalized to two reference genes. Data are shown as the mean ±SD for n=6 (A) and n=3 (B) independent experiments. Asterisks indicate significant differences (\*\**P*<0.01, \*\*\**P*<0.001, \*\*\*\**P*<0.0001; two-tailed *t*-test).

**Supplemental Figure S4. A** Quantification of anthocyanin content in five-day etiolated wild-type (wt) and *tpst-1* seedlings with (+) or without (-) 1 μM CLEL9 in the media. **B** Expression of *CHS*, F3H, *DFR*, *LDOX, UF3GT, UGT78D2, PAP1* and *MYB11* in five-day-old etiolated wt seedlings grown in presence of 1 μM CLEL6 relative to the wt control. Gene expression was normalized to two reference genes. (A,B) Data are shown as the mean ±SD for n=3 independent experiments. Asterisks indicate significant differences (\*\**P*<0.01, \*\*\**P*<0.001, \*\*\*\**P*<0.0001; two-tailed *t*-test).

**Supplemental Figure S5. CLEL9 does not regulate anthocyanin content. A** Quantification of anthocyanin content in five-day-old etiolated wild-type (wt) and *35S:CLEL9* seedlings after 16 h of light exposure. **B** Quantification of anthocyanin content in five-day-old etiolated wt and *35S:CLEL9* seedlings. (A,B) Data are shown as the mean ±SD of n=3 independent experiments.

**Supplemental Figure S6. CLEL6-induced hypocotyl growth depends on RGI receptors. A** Quantification of hypocotyl length of five-day-old etiolated wild-type (wt) and *35S:CLEL6* seedlings. **B** Quantification of hypocotyl length of five-day-old etiolated wt, *tpst-1* and *rgi5x* seedlings grown with (+) or without (−) 1 μM CLEL6 in the media. Asterisks indicate significant differences between the mutants and wt, and between CLEL6-treated and untreated plants. (A,B) Data are shown as the mean ±SD for n=3 independent experiments. Asterisks indicate significant differences (\**P*<0.05, \*\*\*\**P*<0.0001; two-tailed *t*-test).

## Parsed Citations

**Doll, N.M., Royek, S., Fujita, S., Okuda, S., Chamot, S., Stintzi, *A.,* Widiez, T., Hothorn, M., Schaller, A., Geldner, N., and Ingram, G. (2020). A two-way molecular dialogue between embryo and endosperm is required for seed development. Science 367: 431–435.**

Google Scholar: <u>Author Only Title Only Author and Title</u>

**Duek, P.D., and Fankhauser, C. (2005). bHLH class transcription factors take centre stage in phytochrome signalling. Trends in Plant Science 10: 51–54.**

Google Scholar: <u>Author Only Title Only Author and Title</u>

**Fernandez, A., Hilson, P., and Beeckman, T. (2013). GOLVEN peptides as important regulatory signalling molecules of plant development. Journal of Experimental Botany 64: 5263–5268.**

Google Scholar: <u>Author Only Title Only Author and Title</u>

**Ghorbani, S., Hoogewijs, K., Pečenková, T., Fernandez, A., Inzé, A., Eeckhout, D., Kawa, D., Jaeger, G. de, Beeckman, T., Madder, A., van Breusegem, F., and Hilson, P. (2016). The SBT6.1 subtilase processes the GOLVEN1 peptide controlling cell elongation. Journal of Experimental Botany 67: 4877–4887.**

Google Scholar: <u>Author Only Title Only Author and Title</u>

**Igarashi, D., Tsuda, K., and Katagiri, F. (2012). The peptide growth factor, phytosulfokine, attenuates pattern-triggered immunity. The Plant Journal for Cell and Molecular Biology 71: 194–204.**

Google Scholar: <u>Author Only Title Only Author and Title</u>

**Jang, I.-C., Yang, J.-Y., Seo, H.S., and Chua, N.-H. (2005). HFR1 is targeted by COP1 E3 ligase for post-translational proteolysis during phytochrome A signaling. Genes & development 19: 593–602.**

Google Scholar: <u>Author Only Title Only Author and Title</u>

**Jiao, Y., Lau, O.S., and Deng, X.W. (2007). Light-regulated transcriptional networks in higher plants. Nat Rev Genet 8: 217–230.**

Google Scholar: <u>Author Only Title Only Author and Title</u>

**Kang, J., Yu, H., Tian, C., Zhou, W., Li, C., Jiao, Y., and Liu, D. (2014). Suppression of Photosynthetic Gene Expression in Roots Is Required for Sustained Root Growth under Phosphate Deficiency. Plant Physiology 165: 1156–1170.**

Google Scholar: <u>Author Only Title Only Author and Title</u>

**Komori, R., Amano, Y., Ogawa-Ohnishi, M., and Matsubayashi, Y. (2009). Identification of tyrosylprotein sulfotransferase in Arabidopsis. Proceedings of the National Academy of Sciences of the United States of America 106: 15067–15072.**

Google Scholar: <u>Author Only Title Only Author and Title</u>

**Koussevitzky, S., Nott, A., Mockler, T.C., Hong, F., Sachetto-Martins, G., Surpin, M., Lim, J., Mittler, R., and Chory, J. (2007). Signals from chloroplasts converge to regulate nuclear gene expression. Science 316: 715–719.**

Google Scholar: <u>Author Only Title Only Author and Title</u>

**Lee, J., He, K., Stolc, V., Lee, H., Figueroa, P., Gao, Y., Tongprasit, W., Zhao, H., Lee, I., and Deng, X.W. (2007). Analysis of transcription factor HY5 genomic binding sites revealed its hierarchical role in light regulation of development. The Plant Cell 19: 731–749.**

Google Scholar: <u>Author Only Title Only Author and Title</u>

**Leivar, P., Monte, E., Oka, Y., Liu, T., Carle, C., Castillon, A., Huq, E., and Quail, P.H. (2008). Multiple phytochrome-interacting bHLH transcription factors repress premature seedling photomorphogenesis in darkness. Current Biology 18: 1815–1823.**

Google Scholar: <u>Author Only Title Only Author and Title</u>

**Lorrain, S., Allen, T., Duek, P.D., Whitelam, G.C., and Fankhauser, C. (2008). Phytochrome-mediated inhibition of shade avoidance involves degradation of growth-promoting bHLH transcription factors. The Plant Journal 53: 312–323.**

Google Scholar: <u>Author Only Title Only Author and Title</u>

**Maier, A., Schrader, A., Kokkelink, L., Falke, C., Welter, B., Iniesto, E., Rubio, V., Uhrig, J.F., Hülskamp, M., and Hoecker, U. (2013). Light and the E3 ubiquitin ligase COP1/SPA control the protein stability of the MYB transcription factors PAP1 and PAP2 involved in anthocyanin accumulation in Arabidopsis. The Plant Journal 74: 638–651.**

Google Scholar: <u>Author Only Title Only Author and Title</u>

**Martín, G., Leivar, P., Ludevid, D., Tepperman, J.M., Quail, P.H., and Monte, E. (2016). Phytochrome and retrograde signalling pathways converge to antagonistically regulate a light-induced transcriptional network. Nature Communications 7: 11431.**

Google Scholar: <u>Author Only Title Only Author and Title</u>

**Matsuzaki, Y., Ogawa-Ohnishi, M., Mori, A., and Matsubayashi, Y. (2010). Secreted peptide signals required for maintenance of root stem cell niche in Arabidopsis. Science 329: 1065–1067.**

Google Scholar: <u>Author Only Title Only Author and Title</u>

**Meng, L., Buchanan, B.B., Feldman, L.J., and Luan, S. (2012). CLE-like (CLEL) peptides control the pattern of root growth and lateral root development in Arabidopsis. Proceedings of the National Academy of Sciences of the United States of America 109:**

**1760–1765.**

Google Scholar: <u>Author Only Title Only Author and Title</u>

**Monte, E., Al-Sady, B., Leivar, P., and Quail, P.H. (2007). Out of the dark: how the PIFs are unmasking a dual temporal mechanism of phytochrome signalling. Journal of Experimental Botany 58: 3125–3133.**

Google Scholar: <u>Author Only Title Only Author and Title</u>

**Mosher, S., Seybold, H., Rodriguez, P., Stahl, M., Davies, K.A., Dayaratne, S., Morillo, S.A., Wierzba, M., Favery, B., Keller, H., Tax, F.E., and Kemmerling, B. (2013). The tyrosine-sulfated peptide receptors PSKR1 and PSY1R modify the immunity of Arabidopsis to biotrophic and necrotrophic pathogens in an antagonistic manner. The Plant Journal 73: 469–482.**

Google Scholar: <u>Author Only Title Only Author and Title</u>

**Murashige, T., and Skoog, F. (1962). A Revised Medium for Rapid Growth and Bio Assays with Tobacco Tissue Cultures. Physiologia Plantarum 15: 473–497.**

Google Scholar: <u>Author Only Title Only Author and Title</u>

**Nakabayashi, R., Yonekura-Sakakibara, K., Urano, K., Suzuki, M., Yamada, Y., Nishizawa, T., Matsuda, F., Kojima, M., Sakakibara, H., Shinozaki, K., Michael, A.J., Tohge, T., Yamazaki, M., and Saito, K. (2014). Enhancement of oxidative and drought tolerance in Arabidopsis by overaccumulation of antioxidant flavonoids. The Plant Journal 77: 367–379.**

Google Scholar: <u>Author Only Title Only Author and Title</u>

**Nakata, M., Mitsuda, N., Herde, M., Koo, A.J.K., Moreno, J.E., Suzuki, K., Howe, G.A., and Ohme-Takagi, M. (2013). A bHLH-type transcription factor, ABA-INDUCIBLE BHLH-TYPE TRANSCRIPTION FACTOR/JA-ASSOCIATED MYC2-LIKE1, acts as a repressor to negatively regulate jasmonate signaling in arabidopsis. The Plant Cell 25: 1641–1656.**

Google Scholar: <u>Author Only Title Only Author and Title</u>

**Osterlund, M.T., Hardtke, C.S., Wei, N., and Deng, X.W. (2000). Targeted destabilization of HY5 during light-regulated development of Arabidopsis. Nature 405: 462–466.**

Google Scholar: <u>Author Only Title Only Author and Title</u>

**Ou, Y., Lu, X., Zi, Q., Xun, Q., Zhang, J., Wu, Y., Shi, H., Wei, Z., Zhao, B., Zhang, X., He, K., Gou, X., Li, C., and Li, J. (2016). RGF1 INSENSITIVE 1 to 5, a group of LRR receptor-like kinases, are essential for the perception of root meristem growth factor 1 in Arabidopsis thaliana. Cell Research 26: 686–698.**

Google Scholar: <u>Author Only Title Only Author and Title</u>

**Rautengarten, C., Steinhauser, D., Büssis, D., Stintzi, A., Schaller, A., Kopka, J., and Altmann, T. (2005). Inferring hypotheses on functional relationships of genes: Analysis of the Arabidopsis thaliana subtilase gene family. PLoS Computational Biology 1: e40.**

Google Scholar: <u>Author Only Title Only Author and Title</u>

**Reichardt, S., Piepho, H.-P., Stintzi, A., and Schaller, A. (2020). Peptide signaling for drought-induced tomato flower drop. Science 367: 1482–1485.**

Google Scholar: <u>Author Only Title Only Author and Title</u>

**Richter, A.S., Tohge, T., Fernie, A.R., and Grimm, B. (2020). The genomes uncoupled-dependent signalling pathway coordinates plastid biogenesis with the synthesis of anthocyanins. Philosophical Transactions of the Royal Society of London. 375: 20190403.**

Google Scholar: <u>Author Only Title Only Author and Title</u>

**Richter, R., Behringer, C., Müller, I.K., and Schwechheimer, C. (2010). The GATA-type transcription factors GNC and GNL/CGA1 repress gibberellin signaling downstream from DELLA proteins and PHYTOCHROME-INTERACTING FACTORS. Genes and Development 24: 2093–2104.**

Google Scholar: <u>Author Only Title Only Author and Title</u>

**Royek, S., Bayer, M., Pfannstiel, J., Pleiss, J., Ingram, G., Stintzi, A., and Schaller, A. (2022). Processing of a plant peptide hormone precursor facilitated by posttranslational tyrosine sulfation. Proceedings of the National Academy of Sciences of the United States of America 119: e2201195119.**

Google Scholar: <u>Author Only Title Only Author and Title</u>

**Schaller, A., Stintzi, A., Rivas, S., Serrano, I., Chichkova, N.V., Vartapetian, AB., Martínez, D., Guiamét, J.J., Sueldo, D.J., van der Hoorn, R.A.L., Ramírez, V., and Vera, P. (2018). From structure to function - a family portrait of plant subtilases. The New Phytologist 218: 901–915.**

Google Scholar: <u>Author Only Title Only Author and Title</u>

**Shao, L., Shu, Z., Sun, S.-L., Peng, C.-L., Wang, X.-J., and Lin, Z.-F. (2007). Antioxidation of Anthocyanins in Photosynthesis Under High Temperature Stress. Journal of Integrative Plant Biology 49: 1341–1351.**

Google Scholar: <u>Author Only Title Only Author and Title</u>

**Sheerin, D.J., Menon, C., zur Oven-Krockhaus, S., Enderle, B., Zhu, L., Johnen, P., Schleifenbaum, F., Stierhof, Y.-D., Huq, E., and Hiltbrunner, A. (2015). Light-activated phytochrome A and B interact with members of the SPA family to promote photomorphogenesis in Arabidopsis by reorganizing the COP1/SPA complex. The Plant Cell 27: 189–201.**

Google Scholar: <u>Author Only Title Only Author and Title</u>

**Shen, H., Moon, J., and Huq, E. (2005). PIF1 is regulated by light-mediated degradation through the ubiquitin-26S proteasome pathway to optimize photomorphogenesis of seedlings in Arabidopsis. The Plant Journal 44: 1023–1035.**

Google Scholar: <u>Author Only Title Only Author and Title</u>

**Shin, D.H., Choi, M., Kim, K., Bang, G., Cho, M., Choi, S.-B., Choi, G., and Park, Y.-I. (2013). HY5 regulates anthocyanin biosynthesis by inducing the transcriptional activation of the MYB75/PAP1 transcription factor in Arabidopsis. FEBS Letters 587: 1543–1547.**

Google Scholar: <u>Author Only Title Only Author and Title</u>

**Shin, J., Kim, K., Kang, H., Zulfugarov, I.S., Bae, G., Lee, C.-H., Lee, D., and Choi, G. (2009). Phytochromes promote seedling light responses by inhibiting four negatively-acting phytochrome-interacting factors. Proceedings of the National Academy of Sciences of the United States of America 106: 7660–7665.**

Google Scholar: <u>Author Only Title Only Author and Title</u>

**Shin, J., Park, E., and Choi, G. (2007). PIF3 regulates anthocyanin biosynthesis in an HY5-dependent manner with both factors directly binding anthocyanin biosynthetic gene promoters in Arabidopsis. The Plant Journal 49: 981–994.**

Google Scholar: <u>Author Only Title Only Author and Title</u>

**Song, W., Liu, L., Wang, J., Wu, Z., Zhang, H., Tang, J., Lin, G., Wang, Y., Wen, X., Li, W., Han, Z., Guo, H., and Chai, J. (2016). Signature motif-guided identification of receptors for peptide hormones essential for root meristem growth. Cell Research 26: 674–685.**

Google Scholar: <u>Author Only Title Only Author and Title</u>

**Stegmann, M., Zecua-Ramirez, P., Ludwig, C., Lee, H.-S., Peterson, B., Nimchuk, Z.L., Belkhadir, Y., and Hückelhoven, R. (2022). RGI-GOLVEN signaling promotes cell surface immune receptor abundance to regulate plant immunity. EMBO Reports 23: e53281.**

Google Scholar: <u>Author Only Title Only Author and Title</u>

**Stintzi, A., and Schaller, A. (2022). Biogenesis of post-translationally modified peptide signals for plant reproductive development. Current Opinion in Plant Biology 69: 102274.**

Google Scholar: <u>Author Only Title Only Author and Title</u>

**Stracke, R., Ishihara, H., Huep, G., Barsch, A., Mehrtens, F., Niehaus, K., and Weisshaar, B. (2007). Differential regulation of closely related R2R3-MYB transcription factors controls flavonol accumulation in different parts of the Arabidopsis thaliana seedling. The Plant Journal 50: 660–677.**

Google Scholar: <u>Author Only Title Only Author and Title</u>

**Stührwohldt, N., Bühler, E., Sauter, M., and Schaller, A. (2021). Precursor processing by SBT3.8 and phytosulfokine signaling contribute to drought stress tolerance in Arabidopsis. Journal of Experimental Botany.**

Google Scholar: <u>Author Only Title Only Author and Title</u>

**Stührwohldt, N., Scholl, S., Lang, L., Katzenberger, J., Schumacher, K., and Schaller, A. (2020). The biogenesis of CLEL peptides involves several processing events in consecutive compartments of the secretory pathway. eLife 9.**

Google Scholar: <u>Author Only Title Only Author and Title</u>

**Tost, A.S., Kristensen, A., Olsen, L.I., Axelsen, K.B., and Fuglsang, A.T. (2021). The PSY Peptide Family-Expression, Modification and Physiological Implications. Genes 12.**

Google Scholar: <u>Author Only Title Only Author and Title</u>

**Wang, J., Li, H., Han, Z., Zhang, H., Wang, T., Lin, G., Chang, J., Yang, W., and Chai, J. (2015). Allosteric receptor activation by the plant peptide hormone phytosulfokine. Nature 525: 265–268.**

Google Scholar: <u>Author Only Title Only Author and Title</u>

**Wang, X., Zhang, N., Zhang, L., He, Y., Cai, C., Zhou, J., Li, J., and Meng, X. (2021). Perception of the pathogen-induced peptide RGF7 by the receptor-like kinases RGI4 and RGI5 triggers innate immunity in Arabidopsis thaliana. New Phytologist 230: 1110– 1125.**

Google Scholar: <u>Author Only Title Only Author and Title</u>

**Whitford, R., Fernandez, A., Tejos, R., Pérez, A.C., Kleine-Vehn, J., Vanneste, S., Drozdzecki, A., Leitner, J., Abas, L., Aerts, M., Hoogewijs, K., Baster, P., Groodt, R. de, Lin, Y.-C., Storme, V., van de Peer, Y., Beeckman, T., Madder, A., Devreese, B., Luschnig, C., Friml, J., and Hilson, P. (2012). GOLVEN secretory peptides regulate auxin carrier turnover during plant gravitropic responses. Developmental Cell 22: 678–685.**

Google Scholar: <u>Author Only Title Only Author and Title</u>

**Xie, D.-Y., Sharma, S.B., Wright, E., Wang, Z.-Y., and Dixon, R.A. (2006). Metabolic engineering of proanthocyanidins through coexpression of anthocyanidin reductase and the PAP1 MYB transcription factor. The Plant Journal 45: 895–907.**

Google Scholar: <u>Author Only Title Only Author and Title</u>

**Yamada, M., Han, X., and Benfey, P.N. (2020). RGF1 controls root meristem size through ROS signalling. Nature 577: 85–88.**

Google Scholar: <u>Author Only Title Only Author and Title</u>

**Zhang, H., Hu, Z., Lei, C., Zheng, C., Wang, J., Shao, S., Li, X., Xia, X., Cai, X., Zhou, J., Zhou, Y., Yu, J., Foyer, C.H., and Shi, K. (2018). A Plant Phytosulfokine Peptide Initiates Auxin-Dependent Immunity through Cytosolic Ca2+ Signaling in Tomato. The Plant Cell 30: 652–667.**

Google Scholar: <u>Author Only Title Only Author and Title</u>

**Zhao, Z., Shuang, J., Li, Z., Xiao, H., Liu, Y., Wang, T., Wei, Y., Hu, S., Wan, S., and Peng, R. (2021). Identification of the Golden-2-like transcription factors gene family in Gossypium hirsutum. Peer J 9: e12484.**

Google Scholar: <u>Author Only Title Only Author and Title</u>

